# Metabolism of *Lactobacillus* and *Gardnerella vaginalis* in vaginal defined media

**DOI:** 10.1101/2025.04.17.649369

**Authors:** Victoria Horrocks, Charlotte K. Hind, J. Mark Sutton, Rachel M. Tribe, A. James Mason

**Author notes:** **Corresponding Author** A. James Mason, King’s College London, Institute of Pharmaceutical Science, Franklin-Wilkins Building 150 Stamford Street, London, SE1 9NH, UK.

## Abstract

Lactobacilli and *Gardnerella vaginalis* are key bacteria within the vaginal microbiota, significantly impacting its chemical environment. Accurately reproducing vaginal bacteria metabolism *in vitro* is therefore critical in understanding how this may contribute to vaginal health and disease. Complex media such as brain heart infusion (BHI) support the growth of a wide variety of vaginal bacteria but may not accurately reproduce the conditions found *in vivo* leading to over- or under-representation of important metabolic processes. Here we use a ^1^H NMR approach to characterise the growth of *Lactobacillus crispatus, L. jensenii* and a panel of *G. vaginalis* isolates that differ in their metabolic strategy in a vaginal defined media (VDM). We find that both *Lactobacillus* species grow well in VDM, produce much more lactate, and consequently acidify the spent culture far more than when cultured in BHI. In contrast, *G. vaginalis* grows less well in VDM but differences in *Bifidobacterium* shunt and mixed acid fermentation, previously observed in BHI and evidenced by formate production, are preserved. Importantly neither species consume the available glucose but nevertheless conduct carbohydrate fermentation, implicating a preference for glycogen over glucose metabolism with glucose release evidence of glycogen breakdown. While further work is needed to develop media that can support the growth of a wider variety of vaginal bacteria species and recapitulate all features of their metabolism, VDM nevertheless accurately models the key aspects of the chemical environment when *Lactobacillus* dominate, and *G. vaginalis* is prevalent, in the microbiota.

## Introduction

The composition of the vaginal microbiota is increasingly the focus of research that seeks to understand how it contributes to a range of health conditions including bacterial vaginosis (BV), preterm birth, pelvic inflammatory disease and sexually transmitted diseases including HIV.^1,2^ Often, the vaginal microbiota is dominated by one or sometimes two species of *Lactobacillus*.^3–7^ However, more diverse microbiota are also common, with *Gardnerella vaginalis* frequently the most abundant species present.^6^ A diverse vaginal microbiota is sometimes associated with poorer outcomes for the conditions mentioned above.^5–8^ However, this is not predictive and the description of diverse vaginal microbiota as dysbiotic may be inaccurate depending on the cohort being studied.^9^

Although *Lactobacillus* species share many similarities, most notably their ability to produce substantial quantities of lactate from carbohydrate fermentation and consequently acidify their environment, there are also several important differences between them and their prevalence varies according to patient population. For example, in one study *Lactobacillus jensenii* dominance was frequently observed.^10^ However, in our own work, while we also found that *Lactobacillus crispatus, Lactobacillus gasseri* and *Lactobacillus iners* can dominate the vaginal microbiota, *Lactobacillus jensenii*, though prevalent may be less likely to achieve dominance and co-exist with other lactobacilli, notably *L. iners*, or alongside a diverse range of bacteria.^8^ In individuals *L. crispatus* dominance is associated with higher lactate production and lower vaginal pH while absence of *Lactobacillus* dominance is associated with reduced lactate concentrations, higher pH and also more acetate.^8^ Consequently the vaginal chemical environment may vary subtly or substantially according to the microbiota composition and there is a need to understand how and why bacterial metabolism varies in the vagina and the extent to which there is functional redundancy or to which changes in chemical composition can be associated with, or even cause, disease.

Previously we have used a ^1^H NMR metabolomics approach to identify two groupings of *G. vaginalis* isolates which use either mixed acid fermentation or the *Bifidobacterium* shunt (BS).^11^ This is likely to have a significant impact on the vaginal environment as the BS is associated with a relative increase in lactate compared with MAF which is associated with relative increases in acetate, ethanol, and formate. We observed much higher lactate production by *Lactobacillus* species compared with bacterial vaginosis associated bacteria (BVAB) including *G. vaginalis*, but the levels of lactate obtained, and the acidification produced, were lower than is commonly observed in samples obtained from *in vivo*.^11^ In our previous study we used brain heart infusion (BHI), supplemented with serum, to support the growth of both lactobacilli and a panel of BVAB. Since all glucose was consumed from BHI by *Lactobacillus* species this, in addition to the availability of other metabolites, could be a limiting factor for the replication of these two key features. Refinement of the culture conditions is needed to recapitulate the vaginal environment more effectively.

BHI is a complex medium which, when supplemented with serum, contains the required nutrients for growth of most vaginal bacteria species. However, it is not representative of the vaginal environment and others have sought to create a more bespoke culture medium. Simulation of vaginal fluid has been proposed,^12^ however, this media was designed for the purposes of testing pharmaceutical agents rather than supporting microbial growth and contains only glucose and not glycogen. Glycogen and glucose are present as the fermentable substrates in the vaginal environment, reports of glycogen concentration range from 4.4-32 g/l and glucose at 6.4 g/l.^13,14^ High glycogen concentration is associated with *L. crispatus* and *L. jensenii* but not *L. iners*,^15^ and free glycogen is positively correlated with *Lactobacillus* abundance and low vaginal pH both in pre- and post-menopause.^16^ Therefore, any bacterial growth medium which mimics the vaginal environment should contain glycogen.

A defined medium was developed by Geshnizgani and Onderdonk to mimic the fluid present on the surface of vaginal epithelial cells.^17^ *L. acidophilus* and *Staphylococcus epidermidis* were used as representative organisms in the design of this medium to establish growth requirements. When added to the medium, glycogen prolonged survival of *L. acidophilus* and was included at a concentration of 1 g/l in addition to glucose at 10.8 g/l. Other requirements for growth were 2 mg/ml of albumin to represent proteins within the vaginal fluid. While free amino acids have been reported in vaginal secretions, these were not included in the medium. Mucin is also added, this is a glycoprotein produced by epithelial cells which acts as a protective barrier and will change the viscosity of the medium.^18^ Urea is also present in the vaginal environment at a concentration of 49 mg/100 ml and is included in the medium at a concentration of 0.05%. Urea can be broken down by urease to ammonia and CO_2_, the ammonia can be used as a nitrogen source for amino acid synthesis. Tween 20 is included in the medium as a source of long-chain fatty acids which promotes *Lactobacillus* growth. Additional growth factors, such as niacinamide, biotin and choline chloride, which were not detected in vaginal fluid, were nevertheless added from a Kao and Michayluk Vitamin Solution to further promote growth. This defined medium has been used to assess co-culture between *Prevotella bivia* and *Gardnerella vaginalis* or *Peptostreptococcus anaerobius* however the assessment of metabolites was limited as a targeted approach was used.^19,20^ Indeed, no large-scale metabolic characterisation has been conducted to determine the metabolism of key species from the vaginal microbiota in physiologically relevant media. Here therefore we use a ^1^H NMR metabolomics approach to characterise consumption and production of the most abundant small molecule metabolites in VDM by *L. jensenii*, two *L. crispatus* isolates and a panel of *G. vaginalis* isolates whose metabolism we previously characterised in BHI.^11^

## Experimental procedures

### Bacterial culture

*G. vaginalis* and *L. crispatus* and *L. jensenii* strains are as described previously.^11^ All *G. vaginalis* isolates were plated onto CBA (Oxoid, Hampshire, UK) containing 5% defibrinated sheep’s blood (Oxoid) and incubated at 37°C for 48 hours under anaerobic conditions generated using Thermo Scientific™ Oxoid™ AnaeroGen™. *L. crispatus* and *L. jensenii* strains were plated onto MRS agar (Sigma Aldrich) and incubated at 37°C for 48 hours under anaerobic conditions. For initial overnight cultures a 1 µl loop of culture was used to inoculate 5 ml of brain-heart infusion (BHI) media with 5% horse serum and incubated at 37°C for 48 hours under anaerobic conditions without shaking. To generate samples for NMR analysis, 50 µl of overnight culture was added to 5 ml of fresh BHI with 5% horse serum and incubated at 37°C for 48 hours under anaerobic conditions without shaking. For each condition, cultures were grown in triplicate on three separate occasions. At least eight independent replicate samples were desired but fastidious growth led to some samples being discarded to avoid introducing undue variation into the analysis of growth strategies.

### Vaginal defined media

The vaginal defined media (VDM) was prepared as described in Geshnizgani and Onderdonk.^17^ The following components were added to 930 mL of ultrapure water and autoclaved for 15 minutes at 121°C: NaCl (60 mM); KCl (20 mM); K_2_HPO_4_ (10 mM); KH_2_PO_4_ (10 mM); glucose (60 mM); and cysteine.HCl (3 mM). Stock solutions were used to give the indicated final concentrations in the media: glycogen (5% in 20 ml used to give 0.1%); mucin (1.25% in 20 ml, 0.025%); Tween 20 (2% in 10 ml, 0.02%); urea 40% in 1.25 ml, 0.05%); vitamin K1 (0.5% in 0.2 ml, 0.01%); hemin (0.5% in 10 ml, 0.05%); albumin (5% in 40 ml, 0.2%); MgSO_4_ (5% in 5 ml, 0.03%); and NaHCO_3_ (4% in 1 ml, 0.004%). These were then sterilised either through autoclaving (glycogen, mucin, Tween 20, urea, vitamin K1, hemin) or filter sterilisation through a 0.22 µm filter (albumin, MgSO_4_, NaHCO_3_). The following components were obtained as Kao and Michayluk Vitamin Solution (Sigma-Aldrich, K3129).

Prior to inoculation into VDM, all bacteria were first cultured in BHI for 48 hours at 37°C. A 5% inoculum from the starter culture was used to inoculate 5 mL of VDM. Optimal growth of strains was observed after incubation for 168 hours at 37°C under anaerobic conditions. Optical density cut-off of 0.1 at 600 nm was used as a marker of successful growth.

### NMR metabolomics

For preparation of samples to be used in metabolomics bacterial cultures were pelleted by centrifuge at 5000 rpm at 4°C. Supernatant was filtered with 0.22 µm membrane to remove bacterial cells and large debris and were stored at −80°C until use. To aid suppression of the water signal and deuterium lock and act as an internal reference, 60 µl of D_2_O + 3-(trimethylsilyl)propionic-2,2,3,3-d4 acid sodium salt (TSP-d4) was added to 570 µl of filtered supernatant. Sample pH was adjusted using NaOH to within 0.2 pH units of the BHI media control. ^1^H NMR spectra were recorded on Bruker 600 MHz Bruker Avance III NMR spectrometer (Bruker BioSpin, Coventry, United Kingdom) equipped with a 5 mm ^1^H, ^13^C, ^15^N TCI Prodigy Probe and a cooled sample changer with all samples kept at 4 °C. The 1D spectra were acquired under automation at a temperature of 298 K using Carr-Purcell-Meiboom-Gill presaturation (CMPG) pulse sequence (cpmgrp1). The parameters of spectra acquisition are 32 transients, a spectral width of 20.83 ppm and 65,536 datapoints. For assignment of metabolite peaks additional spectra, Total correlation spectroscopy (TOCSY), ^1^H-^13^C heteronuclear single quantum correlation spectroscopy and J-resolved spectroscopy (JRES), were acquired from a pooled sample containing a small volume of all samples. Resonance positions are quoted in ppm with respect to the methyl peak of TSP-d4 at 0.0 ppm.

All spectra were Fourier transformed in Bruker software and adjusted using automatic baseline correction and phasing in Bruker TopSpin 4.1.3. Multiple databases were used for the assignment of metabolites; Chenomx NMR suite software (Chenomx Inc, Canada), Human Metabolome Database (HMDB) and Biological Magnetic Resonance Data Bank (BMRB).^21^ To convert NMR intensity to mM concentration the Chenomx software programme was used, calibrated to the concentration of TSP-d4 present in the sample and adjusted for dilution by D_2_O. NMR raw data (10.6084/m9.figshare.28311782), NMR assignments (10.6084/m9.figshare.28321904) and calculated metabolite concentrations (10.6084/m9.figshare.28292156) are available at Figshare.

## Results

Quantitative aerobic and anaerobic culture has shown that predominant *Lactobacillus* is present at around 10^8^ CFU/ml in the vagina,^22^ while the bacterial cell density of *G. vaginalis* ranges between 10^5^ and 10^8^ CFU/ml in women with BV.^23–25^ As such, to faithfully model both *Lactobacillus* dominant and *G. vaginalis* enriched microbiota, the final cell density of *G. vaginalis* should match, or at least be within one log_10_ of, that of the *Lactobacillus* species. Growth of the three *Lactobacillus* isolates in VDM is robust and is reproducible for *L. crispatus 2* and *L. jensenii* 2 with some variation noted for *L. crispatus* 1 (Fig. 1A). In contrast none of the *G. vaginalis* isolates match the growth of the *Lactobacillus* isolates in VDM, with KC2 and KC3 reaching O.D._600_ of respectively only 0.191 and 0.158 (Fig. 1A). This is in marked contrast to their growth in BHI where, except for KC1, final O.D._600_ always exceeds 1.00 (Fig. 1B). However, although the growth of *G. vaginalis* is significantly lower in VDM than in BHI (*p* < 0.0001) it is within the range that might be expected *in vivo* and hence investigation of the metabolism of both *Lactobacillus* and *G. vaginalis* grown in these conditions is appropriate.

**Figure 1.**
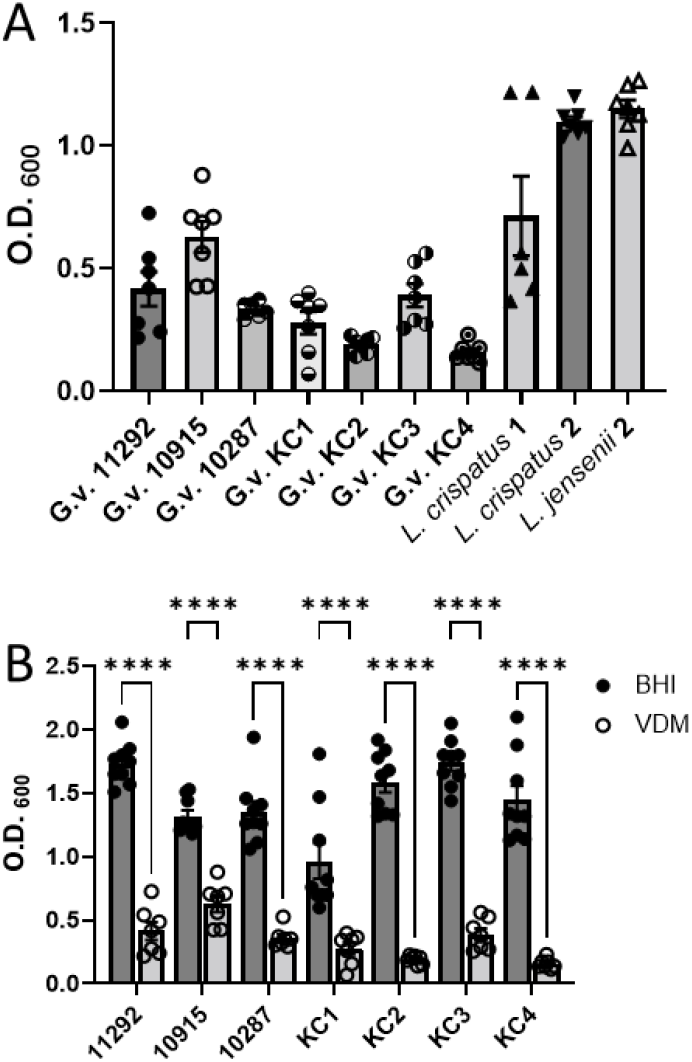
*Gardnerella vaginalis* does not thrive in vaginal defined media. Growth of isolates found within the vaginal microbiota in VDM after 168 hours of incubation (A). Isolates were first cultured in BHI medium for 48 hours and a 5% inoculum was used in the VDM. Comparison of *G. vaginalis* growth in BHI and VDM as determined by Two-way ANOVA with Šídák correction for multiple comparisons, **** *p* < 0.0001. (B). All incubations were carried out at 37°C under anaerobic conditions.

Accordingly, ^1^H NMR metabolomics was used to quantify twenty different metabolites in the spent cultures obtained after growth of either the three *Lactobacillus* or seven *G. vaginalis* isolates (Fig. 2/3; Supp. Fig. 1-3).

**Figure 2.**
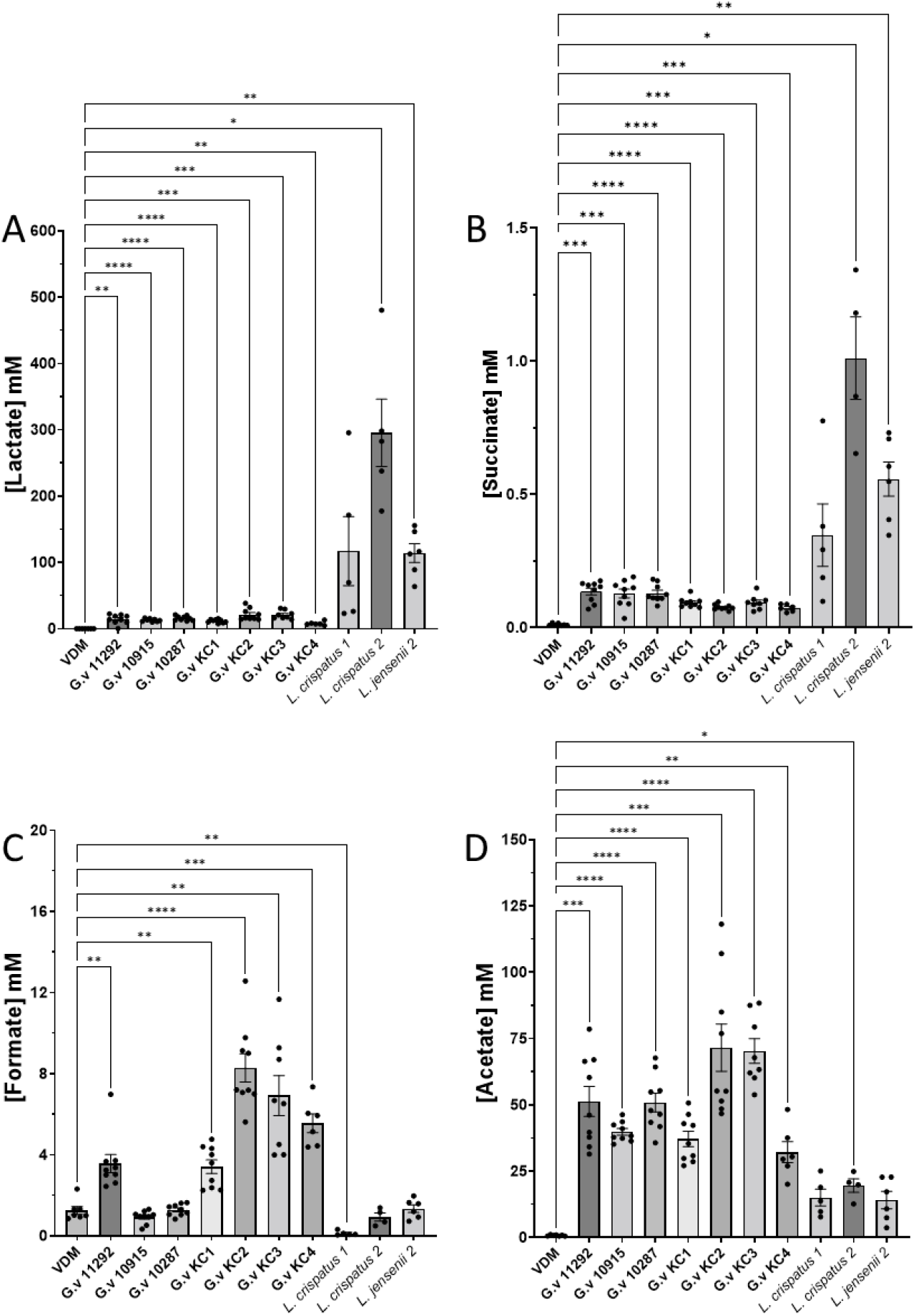
Both *Gardnerella vaginalis and* lactobacilli conduct carbohydrate fermentation in vaginal defined media. Comparison of carbohydrate fermentation products by species found within the vaginal microbiota. ^1^H NMR was used for the identification of metabolites produced from VDM including lactate (A), succinate (B), formate (C) and acetate (D). Log10 transformed data is also presented for lactate (C) Metabolite concentrations were calculated using Chenomx standardised to the concentration of TSP. Statistical test was conducted using a Brown-Forsythe and Welch ANOVA with Dunnett T3 multiple comparisons correction, comparing the concentration produced by isolates to the concentration in fresh VDM media. Only pairwise comparisons where *p* < 0.05 are shown. * *p* < 0.05; ** *p* < 0.01; *** *p* < 0.001; **** *p* < 0.0001.

**Figure 3.**
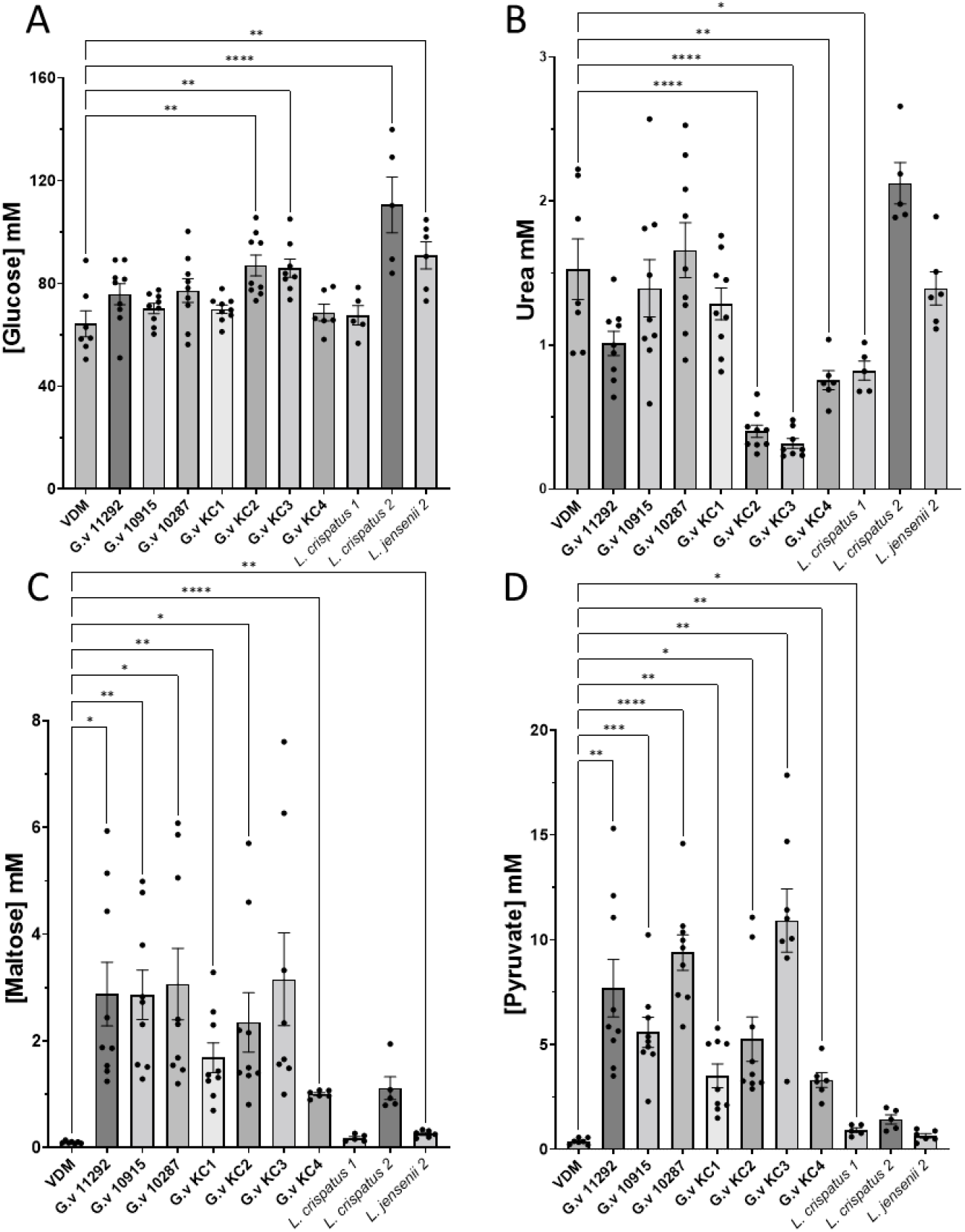
Neither *Gardnerella vaginalis* nor lactobacilli consume glucose from vaginal defined media. Comparison of fermentable substrate metabolism by species found within the vaginal microbiota. ^1^H NMR was used for the identification of metabolites consumed or produced from VDM. Metabolite concentrations were calculated using Chenomx standardised to the concentration of TSP. Statistical test was conducted using a one-way ANOVA (glucose and urea) or a Brown-Forsythe and Welch ANOVA with Dunnett T3 multiple comparisons correction (maltose and pyruvate) comparing the concentration produced by isolates to the concentration in fresh VDM media. Only pairwise comparisons where *p* < 0.05 are shown. * *p* < 0.05; ** *p* < 0.01; *** *p* < 0.001; **** *p* < 0.0001.

### Organic acid production is consistent with carbohydrate fermentation

In BHI both *Lactobacillus* and *G. vaginalis* ferment glucose to produce a range of organic acids, predominantly lactate and acetate, with five of seven *G. vaginalis* strains additionally producing formate via mixed acid fermentation.^11^ In VDM, lactate is produced in substantial quantities by all three *Lactobacillus* strains, ranging from 114 to 295 mM (Fig. 2A), and this is much greater than the circa 40 mM final concentration achieved when the same isolates are cultured in BHI.^11^ The production of lactate by the *G. vaginalis* strains, which ranges between 8 and 22 mM, is significant and, though substantially lower than that observed for the *Lactobacillus* strains, is comparable to that obtained in BHI (7-17 mM) despite a much lower final cell density. The production of succinate follows a similar pattern to that of lactate albeit with around two orders of magnitude difference in concentrations such that while succinate levels are significantly increased in *G. vaginalis* spent cultures this is negligible compared with the production of acetate or lactate (Fig. 2B).

Consistent with the metabolic strategies identified in BHI,^11^ in VDM, formate is again produced by the same five isolates (KC1-4 and NCTC 11292) and not by either of NCTC 10287 and 10915 (Fig. 2C), which are again restricted to using the bifidobacterium shunt, producing only lactate and acetate (Fig. 2D). Spent culture formate concentrations, for the five MAF using *G. vaginalis* strains, range from 3-8 mM and this is much lower than the 26-34 mM obtained in BHI.^11^ Acetate production by *G. vaginalis* ranges from 41-62 mM (Fig. 2D) and, despite the much lower final cell density achieved, this is comparable to that previously achieved in BHI (40-70 mM).^11^ Notably therefore the ratio of acetate to formate produced by the five MAF *G. vaginalis* strains increases from 1.99 ± 0.17 in BHI to 9.98 ± 3.15 in VDM (*p* = 0.0005 by unpaired t test). Acetate is also produced, albeit to a lesser extent, by *Lactobacillus* (*L. crispatus* 1 *p* = 0.0605; *L. jensenii p* = 0.0630). Production ranges from 14-20 mM and this compares with a range of 14-24 mM in BHI,^11^ but this means that the ratio of lactate to acetate production increases from 2.02 ± 0.44 in BHI to 10.32 ± 4.15 in VDM (*p* = 0.0261 by unpaired t test) i.e. *Lactobacillus* increases production of lactate relative to acetate in VDM. *G. vaginalis* also increases production of lactate relative to acetate in VDM such that the ratio of acetate to lactate decreases from 5.82 ± 2.58 in BHI to 3.40 ± 0.34 in VDM (*p* = 0.0302).

Taken together, although qualitatively the production or organic acids by *Lactobacillus* and *G. vaginalis* is the same in VDM as that previously described in BHI there are substantial differences in the absolute and relative amounts of each organic acid. Notably, lactate production by *Lactobacillus* increases substantially in VDM relative to BHI but acetate production is largely unchanged. In contrast, despite relatively limited growth in VDM, *G. vaginalis* nevertheless produces comparable amounts of acetate and lactate to that produced in BHI but, for MAF isolates, much less formate.

### Spent culture and vaginal pH are comparable

Vaginal pH associated with *Lactobacillus* dominance has been found to range between pH 3.6 and 4.5.^4^ Absence of *Lactobacillus* dominance and BV are both associated with an increase in vaginal pH.^4, 26^ The spent culture pH is a function of the production and consumption of acidic or basic metabolites. Notably this includes contributions from the organic acids described above, as well as pyruvate (pK_a_ 2.50) and urea (pK_b_ 13.9) as well as Na HCO_3_ (pK_a_ 6.34) and other ^1^H NMR invisible metabolites such as ammonia. The starting pH of VDM is 6.42 ± 0.07 and all spent cultures are acidified relative to the fresh media (*p* < 0.0001). For *Lactobacillus* the spent culture pH all reaches substantially more acidic pH than in BHI (range 6.03-6.56);^11^ *L. crispatus* 1 4.09 ± 0.26, *L. crispatus* 2 3.46 ± 0.05, *L. jensenii* 3.93 ± 0.05. These values are consistent with what is expected from a *Lactobacillus* dominated microbial community and likely reflect the substantial production of lactate by these isolates (pK_a_ 3.86). In contrast the pH of the *G. vaginalis* spent cultures range from 4.80 to 5.69 and there is no correlation between spent culture pH and the O.D._600_. These values are nevertheless comparable with those previously achieved in BHI, where the spent culture pH ranged from 5.22-5.80,^11^ and are consistent with vaginal pH (pH 5.3) observed in the absence of *Lactobacillus* dominance (and high *G. vaginalis* prevalence).^4^

### Carbohydrate fermentation is not sustained by glucose in VDM

In BHI both *G. vaginalis* and *Lactobacillus* consume all glucose in the media. In VDM, glucose is added at a final concentration of 60 mM and quantification by NMR is consistent with this, with a concentration of 64.3 ± 5.0 determined by comparison with the internal standard (Fig. 3A). Interestingly culture of neither the *G. vaginalis* nor the lactobacilli leads to any net consumption of glucose from the VDM. Indeed, and in stark contrast to what is observed for culture of these strains in BHI, there is a modest release of glucose for cultures of two *G. vaginalis* strains (KC2 and KC3) as well as *L. crispatus* 2 and *L. jensenii* 2 with glucose concentrations in spent culture increasing by between 21.6 mM (KC3) and 46.28 mM (*L. crispatus* 2). Maltose is a disaccharide formed from two glucose monomers and is also a degradation product of glycogenolysis. Maltose is essentially absent from fresh VDM but significant production is detected in all *G. vaginalis* spent cultures as well as in that of *L. crispatus* 2 and *L. jensenii* 2 (Fig. 3C).

When cultured in BHI the five MAF *G. vaginalis* strains additionally consumed pyruvate but spent cultures from the two BS strains had slightly higher pyruvate concentrations than the fresh media.^11^ Since VDM does not contain pyruvate it cannot be consumed, but interestingly modest increases in pyruvate (pK_a_ 2.50) are observed for all seven isolates (Fig. 3D). Modest pyruvate production is also observed for both *L. crispatus* isolates (*L. crispatus* 2 *p* = 0.058).

Genomics analysis of three *G. vaginalis* strains suggests this species is capable of using both amino acids and ammonia as nitrogen sources but not urea,^27^ though others have identified ureases in some isolates.^28^ In our previous work however we did not observe any substantial consumption of amino acids from BHI and in some cases we instead observed modest production.^11^ In VDM we observe consumption of urea by *G. vaginalis*, but only by three of seven strains indicating this is not a generic nitrogen source for the species (Fig. 3B). Interestingly, the two strains that consume the most urea (KC2 and KC3) also have the highest spent culture pH (respectively 5.46 and 5.69) and hence spent culture pH may be associated with production of ammonia though this is invisible to ^1^H NMR. Amino acids are either absent or present in negligible quantities in fresh VDM but the VDM spent culture amino acid content was enriched after culture of all seven *G. vaginalis* strains either as a result of *de novo* synthesis or degradation of albumin and/or mucin (Supp. Fig. 1-3). *L. crispatus* 1 spent culture was modestly depleted of urea (Fig. 3B), but that of the other two *lactobacillus* strains was unaffected. *Lactobacillus* conditioned media was also enriched with amino acids, as observed for *G. vaginalis* (Supp. Fig. 1-3).

## Discussion

Here we use NMR metabolomics to generate a metabolic profile of spent culture media after growth of three *Lactobacillus* and seven *G. vaginalis* isolates in a defined medium designed to mimic vaginal secretions. We aim to understand whether and to what extent a defined medium may provide a more faithful model of the chemical environment that sustains and is a product of microbial growth reflecting both *Lactobacillus* dominated microbiota and more diverse communities in which *G. vaginalis* is often prevalent. Specifically we compare the metabolism of these ten strains with their metabolism when cultured under identical conditions but in BHI.^11^ Whereas BHI is a complex medium, which consists of beef heart and calf brains in addition to peptone and glucose, designed for growth of fastidious bacteria, and does therefore not reflect the vaginal environment, VDM contains both glucose and glycogen as fermentable substrates and contains albumin and mucin to represent protein components within vaginal fluid. We find three key differences: 1) both *Lactobacillus* and *G. vaginalis* eschew glucose, most likely, in favour of glycogen consumption; 2) growth of *G. vaginalis* in VDM is not as robust as in BHI; but 3) despite some aspects that can be improved, the chemical environment of the respective VDM spent cultures nevertheless better reflects the chemical environment of both *Lactobacillus* dominant and *G. vaginalis* enriched microbiota.

### Do both Lactobacillus and G. vaginalis prefer glycogen to glucose?

Since glycogen is a large polymer it cannot be quantified using liquid-state ^1^H NMR, with the technique sensitive only to low molecular weight metabolites containing non- or sufficiently slowly-exchangeable hydrogen. As such the method is limited to quantifying the products of fermentation and the amount of glucose which is the only other carbon source available in substantial quantities. Nevertheless, since there was no net consumption of glucose after culturing either the *Lactobacillus* or *G. vaginalis*, and indeed for some isolates there is a significant increase in either glucose or maltose – both products of glycogen degradation - and with the clear evidence of fermentation for all isolates we can conclude that glycogen is being directly consumed in preference to the 60 mM glucose present in VDM.

While it has been known for a long time that vaginal glycogen sustains *Lactobacillus*, with free glycogen correlating with *Lactobacillus*,^16^ our understanding of how this happens is incomplete. Vaginal epithelial cells produce glycogen as a major carbon source,^29^ but it was originally thought that vaginal *Lactobacillus* were not able to ferment glycogen,^30^ and instead *α*-amylase in the vaginal environment is required to release digestible sugars such as maltose for utilisation by *Lactobacillus* species.^31^ More recently, evidence has emerged that in fact most *Lactobacillus* are indeed capable of glycogen degradation,^32^ with comparative genomics of 28 *L. crispatus* strains identifying genes related to glycogen metabolism.^32^ Of these strains, 25 were tested in glucose-free medium supplemented with glycogen, and all but one strain were able to grow in this medium although six strains exhibited poor growth. Further, recent work has shown that both glycogen availability and pH affects the growth of *L. crispatus, L. jensenii, L. gasseri* and also *G. vaginalis* in medium simulating vaginal fluid (MSVF), with only *L. gassseri* able to survive in MSVF without glycogen.^33^

As such, *Lactobacillus* can be generally considered capable of using glycogen as its primary carbon source. However, since this is not universal there may be scope for species- or strain-specific variation in the ability to consume glycogen to affect microbiota composition and, as discussed below, the overall chemical environment in the vagina. Further, while it is becoming clear that most *Lactobacillus* can and will directly catabolise glycogen, the present observation that three *Lactobacillus* strains preferentially consume glycogen and not glucose, suggests further research will be needed to establish how different species or strains vary in sensitivity to catabolite repression.

Similarly, both genomic analysis of *G. vaginalis*, which suggested all strains are capable of catabolising both glycogen and glucose,^27^ and the above-mentioned study performed in MSVF, indicate that *G. vaginalis* too can directly catabolise glycogen, with the present study indicating that this also happens in preference to glucose consumption. Glycogen therefore will be an important carbon source in both *Lactobacillus* dominated and more diverse microbiota and the presence of glucose may be related to increased glycogen degradation. Both we and others have found an increase of glucose concentration to be associated with preterm birth,^8, 34^ while NMR metabolomics in vaginal samples from women who have BV, identified maltose as an indicator of a eubiotic microbiota and an increased vaginal glucose concentration in dysbiosis.^35^ Further, a meta-RNA-seq analysis found that there were an increase in transcripts associated with glycogen metabolism in BV-positive samples.^36^ BV is associated with significant bacterial overgrowth which is 100-1000 times greater than in the normal flora.^37^ The increase in glucose observed for both BV and sPTB could therefore be a result of increased glycogen breakdown both from invading species as well as a greater number of bacteria present.

### VDM may need further modification to sustain the growth of a more diverse selection of vaginal bacteria and to model other aspects of its chemical environment

VDM contains physiologically accurate concentrations of metabolites, including glycogen, urea and mucin, which are not present in BHI. On the other hand, absent from fresh VDM are amino acids which are identified in both CVF and BHI,^8, 11^ consistent with mass spectrometry metabolomics of vaginal fluid.^38^ These differences may impact both the ability of *Lactobacillus* and *G. vaginalis* to grow as well as the metabolic strategy that they adopt and consequently the ability of VDM to accurately model differing vaginal chemical environments. Compared with BHI the growth of *G. vaginalis* in VDM is limited, with 168 hours of incubation required to at least reach the minimum growth cut off and even then a significantly lower final cell density is achieved at 168 hours compared with BHI at 48 hours.

In the development of VDM *L. acidophilus* was used to test whether the medium supported growth therefore it is unsurprising that the *Lactobacillus* species exhibited superior growth.^17^ However, the absence of arginine, asparagine, isoleucine, lysine, tryptophan and uracil from VDM is an obstacle to understanding whether more diversity in metabolism exists among *Lactobacillus* or *G. vaginalis*. Notably in BHI, *L. crispatus* 1 consumes a wider range of metabolites than *L. crispatus 2* which is restricted to consuming glucose and pyruvate.^11^ *L. crispatus* 1 additionally consumes asparagine, isoleucine, lysine, tryptophan and uracil and both it and *L. jensenii* 2, uniquely among strains tested, consume arginine.^11^ Ornithine is frequently generated from arginine and its production was detected in BHI for both *L. jensenii* and *L. crispatus* 1. Putrescine, its decarboxylation product, was also produced by *L. crispatus* 1.^11^ No ornithine or putrescine are produced by either isolate in VDM and, as such, the absence of arginine from VDM in particular may limit the ability to model biogenic amine production, associated with BV,^39^ by *Lactobacillus* or other species. The absence of uracil from VDM may also be critical since it is consumed from BHI by all *G. vaginalis* and *L. crispatus* 1,^11^ is negatively associated with *L. crispatus* and *L. jensenii* in patients,^40^ and for other bacterial species this is important in anaerobic growth.^41, 42^

While the absence of these amino acids and uracil is a limitation of VDM, a potential advantage is the presence of 0.05% urea as a nitrogen source. In patients, urea is positively correlated with both *L. crispatus* and *L. jensenii*,^40^ decreased in BV,^43^ and yet genome sequencing of three *G. vaginalis* strains indicated that none of these were likely capable of using urea.^27^ The present study clarifies this situation by indicating that while urea consumption by *G. vaginalis* is possible, it is far from universal. Furthermore, while *Lactobacillus* might generally be considered not to consume urea some isolates, represented by *L. crispatus* 1, may have more diverse metabolic strategies. Intra-species variation in urea consumption and consequent ammonia production could be a further contribution to variation in pH beyond the production of organic acids.

### VDM spent culture nevertheless models key aspects of the vaginal chemical environment

The reliability of the two media in reproducing the chemical environment of the vagina may be most critically assessed by considering the production of organic acids which largely determine one of its key features – its acidic pH. The concentration of lactate in the *Lactobacillus* dominated vagina is, on average, 120 mM,^44^ and this compares well with production of lactate by the *Lactobacillus* isolates in VDM (114 – 295 mM). In contrast growth of *Lactobacillus* in BHI produced around a third as much lactate and the spent cultures were not acidified to the same extent. In these respects, VDM substantially outperforms BHI and the origin of this difference should be investigated further. Notably, while the range of fermentation products are qualitatively the same in BHI and VDM, in the latter the increase in lactate likely comes at the expense of acetate production and raises a question as to whether and how this results from a shift from glucose to glycogen consumption? However, since strong spent culture acidification has been reported for *Lactobacillus* culture in de Man, Rogosa and Sharpe (MRS) broth there may be other factors beyond the availability of glycogen that affect lactate production and/or spent culture pH.

Lactate concentrations in the vagina decrease substantially during BV with a value of 20 mM lactate reported.^44^ This is consistent with the range of lactate concentrations observed here for *G. vaginalis* cultured in VDM. Although this relatively low amount of lactate could be attributed to poor growth of *G. vaginalis* in VDM, final acetate concentrations nevertheless ranged between 41 – 62 mM post culture (for *Lactobacillus* this ranges from 14 – 20 mM). Vaginal acetate levels have been found to be higher (10.5 vs 9.0 mM) for women who deliver preterm compared with term,^46^ and a threshold of 15 mM has been used for positive identification of BV.^47^ Therefore, although the growth of *G. vaginalis* in VDM is significantly less than that achievable in BHI, VDM nevertheless supports growth that approximately matches that observed *in vivo* and, importantly this is sufficient to reproduce the expected chemical environment. Again, a shift in the ratio of lactate to acetate production is observed and this warrants further investigation to determine whether and how this is determined by glycogen availability. Also, acidification of MRS broth exceeds that observed in BHI, albeit to a lesser extent,^45^ so other factors may again impact this behaviour.

In BHI two distinct metabolic pathways for *G. vaginalis* were observed with isolates using either the *Bifidobacterium* shunt or mixed acid fermentation.^11^ The production of formate is therefore a distinctive feature of five of the seven *G. vaginalis* strains. In VDM, although the MAF strains continue to produce formate, a substantial increase in acetate production is observed in the MAF strains. This is achieved at the expense of formate production such that now the difference in metabolism between the BS and MAF strains is less substantial while there is no significant difference in production of acetate and lactate between BS and MAF strains; lactate production was higher for BS strains in BHI.^11^

## Conclusion

The culture of *Lactobacillus* and *G. vaginalis* in vaginal defined media: 1) reveals their likely preference for glycogen over glucose as a carbon source; 2) identifies release of glucose from glycogen thus providing further context for biomarker interpretation; 3) demonstrates intra-species variation in urea consumption and; 4) models the chemical environment created by both *Lactobacillus* dominated microbiota and situations where *G. vaginalis* is more prevalent better than brain heart infusion. Modification of VDM may however be needed to sustain better growth of *G. vaginalis* and to better model more diverse metabolic strategies and model the production of biogenic amines.

## ASSOCIATED CONTENT

### Supporting Information

Further comparison of metabolites produced by *Lactobacillus* and *G. vaginalis* is provided as Supplementary Figures S1-S3.

### Data availability

Raw NMR data files, assignments and concentrations are available from Figshare.

## AUTHOR INFORMATION

### CRediT author statement

**Victoria Horrocks:** Conceptualization, Data curation, Formal analysis, Investigation, Methodology, Writing – original draft. **Charlotte Hind:** Conceptualization, Investigation, Methodology, Resources, Supervision. **Mark Sutton:** Conceptualization, Funding acquisition, Project administration, Resources, Supervision. **Rachel Tribe:** Conceptualization, Funding acquisition, Resources, Supervision, Writing – review & editing: **James Mason:** Conceptualization, Data curation, Formal analysis, Funding acquisition, Project administration, Supervision, Writing – original draft, Writing – review & editing.

### Notes

The authors declare no competing interests.

## Acknowledgment

NMR experiments described in this paper were carried out using the facilities of the Centre for Biomolecular Spectroscopy, King’s College London using instruments acquired with a Multi-user Equipment Grant from the Wellcome Trust and an Infrastructure Grant from the British Heart Foundation. We thank Dr Andrew Atkinson, Dr Adrien Le Guennec and Dr James Jarvis for assistance with liquid-state NMR experiments performed at KCL. VH was supported by a King’s College London iCASE award, affiliated to the London Interdisciplinary Doctoral Programme (LIDo), and Public Health England. Funding for the INSGHT cohort providing swabs was provided from Tommy’s Charity (no. 1060508); NIHR Biomedical Research Centre (BRC) based at Guy’s and St. Thomas’ National Health Service Foundation Trust, Rosetrees Trust (charity no. 298582) (M303-CD1) and Borne Foundation (charity no. 1167073). The views expressed are those of the author(s) and not necessarily those of the NHS, the NIHR, or the Department of Health and Social Care. We thank Collette Allen at SDH for providing patient swabs

## Supplementary Materials

**Supplementary Figure 1. Production of alkyl sidechain amino acids by *Gardnerella vaginalis* and lactobacilli from vaginal defined media.** Comparison of amino acid production by species found within the vaginal microbiota. ^1^H NMR was used for the identification of metabolites produced from VDM including isoleucine (A), alanine (B), leucine (C) and valine (D). Metabolite concentrations were calculated using Chenomx standardised to the concentration of TSP. Statistical test was conducted using a Brown-Forsythe and Welch ANOVA with Dunnett T3 multiple comparisons correction (maltose and pyruvate) comparing the concentration produced by isolates to the (almost zero) concentration in fresh VDM media. Only pairwise comparisons where *p* < 0.05 are shown. * *p* < 0.05; ** *p* < 0.01; *** *p* < 0.001; **** *p* < 0.0001.

**Supplementary Figure 2. Production of polar amino acids by *Gardnerella vaginalis* and lactobacilli from vaginal defined media.** Comparison of amino acid production by species found within the vaginal microbiota. ^1^H NMR was used for the identification of metabolites produced from VDM including lysine (A), glutamate (B), asparagine (C) and aspartate (D). Metabolite concentrations were calculated using Chenomx standardised to the concentration of TSP. Statistical test was conducted using a one-way ANOVA (asparagine and aspartate) or a Brown-Forsythe and Welch ANOVA with Dunnett T3 multiple comparisons correction (lysine and glutamate) comparing the concentration produced by isolates to the concentration in fresh VDM media. Only pairwise comparisons where *p* < 0.05 are shown. * *p* < 0.05; ** *p* < 0.01; *** *p* < 0.001; **** *p* < 0.0001.

**Supplementary Figure 3. Production of methionine and aromatic amino acids by *Gardnerella vaginalis* and lactobacilli from vaginal defined media.** Comparison of production by species found within the vaginal microbiota. ^1^H NMR was used for the identification of metabolites produced from VDM including methionine (A), tryptophan (B), tyrosine (C) and phenylalanine (D). Metabolite concentrations were calculated using Chenomx standardised to the concentration of TSP. Statistical test was conducted using a Brown-Forsythe and Welch ANOVA with Dunnett T3 multiple comparisons correction comparing the concentration produced by isolates to the concentration in fresh VDM media. Only pairwise comparisons where *p* < 0.05 are shown. * *p* < 0.05; ** *p* < 0.01; *** *p* < 0.001; **** *p* < 0.0001.

