## Supplementary figures for "Metabolism of *Lactobacillus* and *Gardnerella vaginalis* in vaginal defined media"

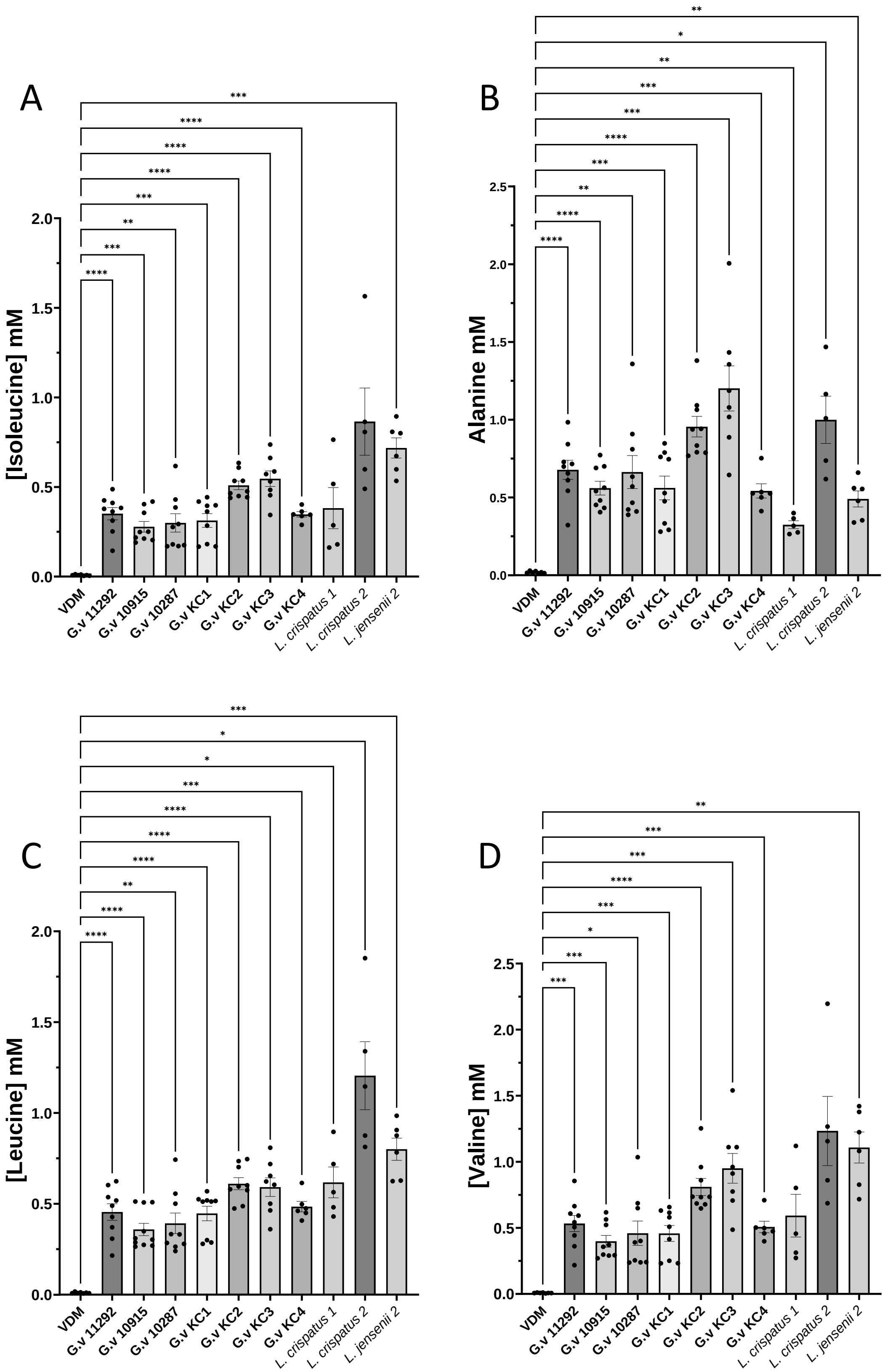


**Supplementary Figure 1. Production of alkyl sidechain amino acids by *Gardnerella vaginalis* and lactobacilli from vaginal defined media.** Comparison of amino acid production by species found within the vaginal microbiota. $^{1}$H NMR was used for the identification of metabolites produced from VDM including isoleucine (A), alanine (B), leucine (C) and valine (D). Metabolite concentrations were calculated using Chenomx standardised to the concentration of TSP. Statistical test was conducted using a Brown-Forsythe and Welch ANOVA with Dunnett T3 multiple comparisons correction (maltose and pyruvate) comparing the concentration produced by isolates to the (almost zero) concentration in fresh VDM media. Only pairwise comparisons where $p < 0.05$ are shown. * $p < 0.05$; ** $p < 0.01$; *** $p < 0.001$; **** $p < 0.0001$.

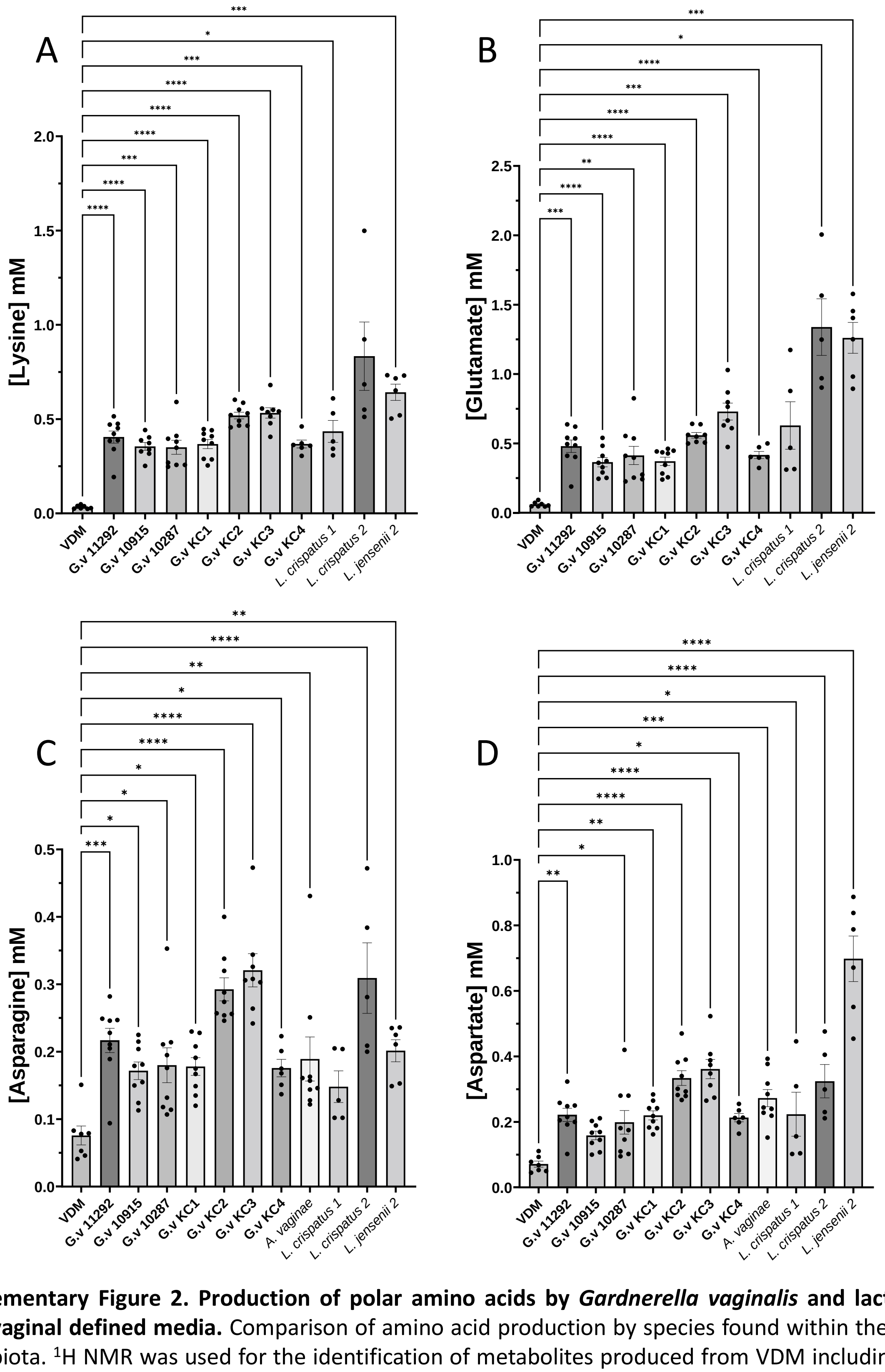


**Supplementary Figure 2. Production of polar amino acids by *Gardnerella vaginalis* and lactobacilli from vaginal defined media.** Comparison of amino acid production by species found within the vaginal microbiota. $^{1}$H NMR was used for the identification of metabolites produced from VDM including lysine (A), glutamate (B), asparagine (C) and aspartate (D). Metabolite concentrations were calculated using Chenomx standardised to the concentration of TSP. Statistical test was conducted using a one-way ANOVA (asparagine and aspartate) or a Brown-Forsythe and Welch ANOVA with Dunnett T3 multiple comparisons correction (lysine and glutamate) comparing the concentration produced by isolates to the concentration in fresh VDM media. Only pairwise comparisons where $p < 0.05$ are shown. * $p < 0.05$; ** $p < 0.01$; *** $p < 0.001$; **** $p < 0.0001$.

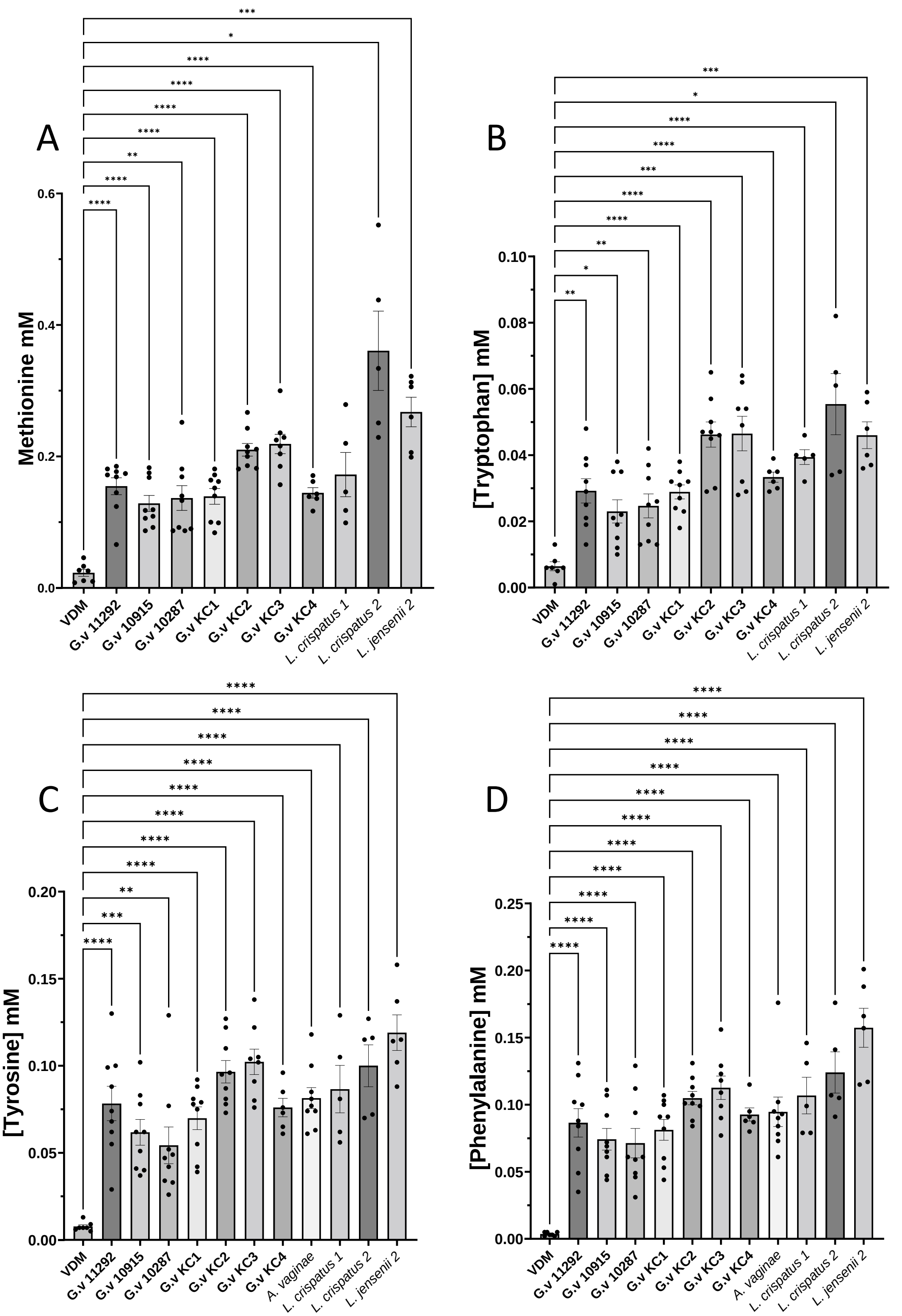


**Supplementary Figure 3. Production of methionine and aromatic amino acids by *Gardnerella vaginalis* and lactobacilli from vaginal defined media.** Comparison of production by species found within the vaginal microbiota. $^1$H NMR was used for the identification of metabolites produced from VDM including methionine (A), tryptophan (B), tyrosine (C) and phenylalanine (D). Metabolite concentrations were calculated using Chenomx standardised to the concentration of TSP. Statistical test was conducted using a Brown-Forsythe and Welch ANOVA with Dunnett T3 multiple comparisons correction comparing the concentration produced by isolates to the concentration in fresh VDM media. Only pairwise comparisons where $p < 0.05$ are shown. * $p < 0.05$; ** $p < 0.01$; *** $p < 0.001$; **** $p < 0.0001$.
